# scTenifoldNet: a machine learning workflow for constructing and comparing transcriptome-wide gene regulatory networks from single-cell data

**DOI:** 10.1101/2020.02.12.931469

**Authors:** Daniel Osorio, Yan Zhong, Guanxun Li, Jianhua Z. Huang, James J. Cai

## Abstract

Constructing and comparing gene regulatory networks (GRNs) from single-cell RNA sequencing (scRNAseq) data has the potential to reveal critical components in the underlying regulatory networks regulating different cellular transcriptional activities. Here, we present a robust and powerful machine learning workflow—scTenifoldNet—for comparative GRN analysis of single cells. The scTenifoldNet workflow, consisting of principal component regression, low-rank tensor approximation, and manifold alignment, constructs and compares transcriptome-wide single-cell GRNs (scGRNs) from different samples to identify gene expression signatures shifting with cellular activity changes such as those associated with pathophysiological processes and responses to environmental perturbations. We used simulated data to benchmark scTenifoldNet’s performance, and then applied scTenifoldNet to several real data sets. In real-data applications, scTenifoldNet identified highly specific changes in gene regulation in response to acute morphine treatment, an antibody anticancer drug, gene knockout, double-stranded RNA stimulus, and amyloid-beta plaques in various types of mouse and human cells. We anticipate that scTenifoldNet can help achieve breakthroughs through constructing and comparing scGRNs in poorly characterized biological systems, by deciphering the full cellular and molecular complexity of the data.

**Highlights:**

- scTenifoldNet is a machine learning workflow built upon principal component regression, low-rank tensor approximation, and manifold alignment
- scTenifoldNet uses single-cell RNA sequencing (scRNAseq) data to construct single-cell gene regulatory networks (scGRNs)
- scTenifoldNet compares scGRNs of different samples to identify differentially regulated genes
- Real-data applications demonstrate that scTenifoldNet accurately detects specific signatures of gene expression relevant to the cellular systems tested.

**Short abstract:** We present scTenifoldNet—a machine learning workflow built upon principal component regression, low-rank tensor approximation, and manifold alignment—for constructing and comparing single-cell gene regulatory networks (scGRNs) using data from single-cell RNA sequencing (scRNAseq). scTenifoldNet reveals regulatory changes in gene expression between samples by comparing the constructed scGRNs. With real data, scTenifoldNet identifies specific gene expression programs associated with different biological processes, providing critical insights into the underlying mechanism of regulatory networks governing cellular transcriptional activities.

## Main

A gene regulatory network (GRN) is a graph depicting the intricate interactions between transcription factors (TFs), associated proteins and their target genes, reflecting the physiological condition of the cells in question. The analysis of GRNs promotes the interpretation of cell states, cell functions, and regulatory mechanisms that underlie the dynamics of cell behaviors. Multiple methods have been developed to build GRNs from data of gene expression [1–4]. It is important to compare GRNs constructed using data sets from different samples because the comparison may reveal regulatory mechanisms leading to transcriptomic changes. In particular, the comparison results may help understand what is the most significant shift in regulatory mechanisms between samples, as well as how genetic and environmental signals are integrated to regulate a cell population’s physiological responses and how cell behavior is affected by various perturbations. All of these are key questions in the study of the functional participation of given GRNs. Despite the critical importance of comparative GRN analysis, relatively few methods have been established to compare GRNs [5].

Single-cell RNAseq (scRNAseq) technology has been revolutionizing the biomedical sciences in recent years. New research provides an unparalleled degree of precision to analyze transcriptional regulation, cell history, and cell interactions with rich knowledge. It transforms previous entire tissue-based assays into transcriptomic single-cell measurements and greatly enhances our understanding of cell development, homeostasis and disease. Current scRNAseq systems (e.g., 10x Genomics) can profile transcriptomes for thousands of cells per experiment. This sheer number of measured cells can be leveraged to construct GRNs. Advanced computational methods can facilitate such an effort to reach unprecedented resolution and accuracy, revealing the network state of given cells [6–8]. Furthermore, comparative analyses among GRNs of different samples will be extremely powerful in revealing fundamental changes in regulatory networks and unraveling the transcriptional programs that govern the behaviors of cells. Since our ability to generate scRNAseq data has outpaced our ability to extract information from it, there is a clear need to develop effective computational algorithms and novel statistical methods for analyzing and exploiting information embedded within GRNs [9].

Constructing single-cell GRNs (scGRNs) using data from scRNAseq and then effectively comparing constructed scGRNs present great analytical challenges [9, 10]. A meaningful comparison of scGRNs first requires a robust construction of GRN from scRNAseq data. Comparing scGRNs built via an unstable solution would cause misleading results and inappropriate conclusions. The vast number of different cellular states in a sample, technical and biological noise, as well as the sparsity of scRNAseq data, complicate the process of scGRN construction. Often, the expression of a gene is governed by stochastic processes and also influenced by transcriptional activities of many other genes. Thus, it is difficult to tease out subtle signals and infer true connections between genes. Furthermore, a direct comparison between two scGRNs is difficult—e.g., comparing each edge of the graph between scGRNs would be ill-powered when scGRNs involve thousands of genes. Taken together, the key challenge in conducting comparative scGRN analysis is to extract meaningful information from noisy and sparse scRNAseq data, since the information is deeply embedded in the differences between highly complex scGRNs of two samples.

In this paper, we introduce a workflow for constructing and comparing scGRNs using data from the scRNAseq of different samples. The workflow, which we call scTenifoldNet, is built upon several machine learning algorithms, including principal component regression, low-rank tensor approximation, and manifold alignment. Through several examples, we show that scTenifoldNet is a sensitive tool to detect specific changes in gene expression signatures and the regulatory network rewiring events. scTenifoldNet inputs are a pair of matrices of scRNAseq expression from two different samples. For instance, one sample may come from a healthy donor and the other from a diseased donor. In scTenifoldNet, the two input expression matrices are simultaneously processed through a multistep procedure. The final output is a list of ranked genes sorted by their significance, assessed by a specifically designed, differential regulation (DR) test. The ranked gene list can be used to perform functional enrichment analysis to detect the enriched molecular functions and involved biological processes. The constructed scGRN can also be used for the identification of functionally significant modules, i.e., subsets of tightly regulated genes.

scTenifoldNet is an innovative method in terms of its scGRN comparison function. We are not aware of any prior work using a similar design to achieve the same analytical goal. scTenifoldNet overcomes several technological challenges in implementing an effective and efficient scGRN comparison method. Here, we first benchmarked and demonstrated the utility of scTenifoldNet across synthetic data sets and then applied scTenifoldNet to real data sets. Our real data analyses showed scTenifoldNet’s power in identifying significant genes and network modules whose regulatory patterns are shifting greatly between samples. Some of these findings have not been reported in the respective original studies, in which the data sets were generated.

## Results

### The scTenifoldNet architecture

To enable comparative scGRN analysis in a robust and scalable manner, we base our method on a series of machine learning methods. A key challenge of our comparative analysis is to extract meaningful differences in regulatory relationships between two samples from noisy and sparse data. Specifically, we seek to contrast scGRNs constructed from different scRNAseq expression matrices. **Fig. 1** shows the main components of scTenifoldNet architecture. The whole workflow contains five key steps: subsampling cells, constructing multilayer scGRNs, denoising, manifold alignment, and differential regulation (DR) test. In order to produce biologically meaningful results, we made dedicated design decisions for the task in each of these steps. Next, we briefly describe the numerical methods implemented in scTenifoldNet. More technical details are presented in **Methods**.

**Fig. 1.**
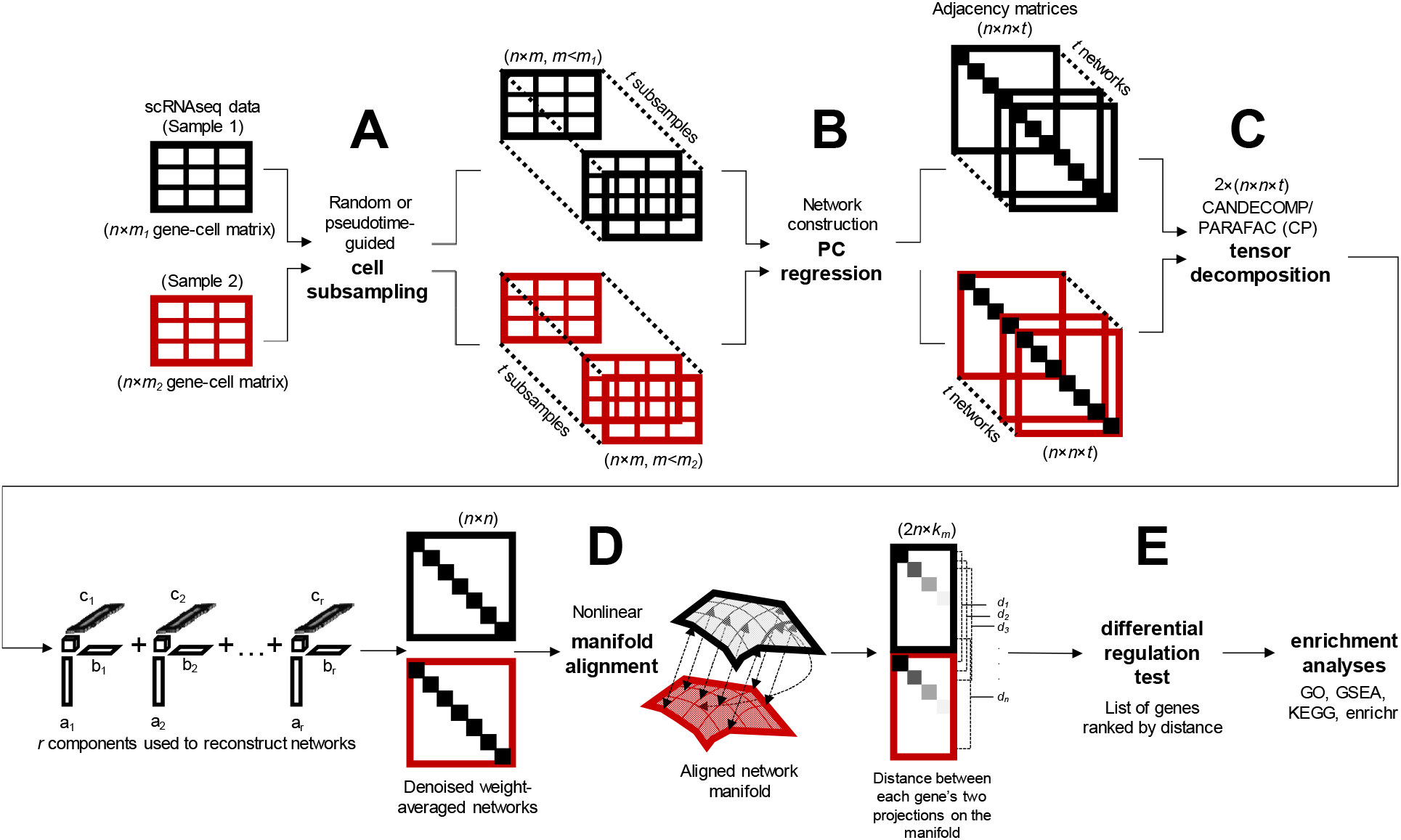
Overview of the scTenifoldNet workflow. scTenifoldNet is a machine learning framework that uses a comparative network approach with scRNAseq data to identify regulatory changes between samples. scTenifoldNet is composed of five major steps. (**A**) *Cell subsampling*. scTenifoldNet starts with subsampling cells in the scRNAseq expression matrices. When two samples are analyzed, each of the two samples is subsampled either randomly or following a pseudotime trajectory of cells. The subsampling is repeated multiple times to create a series of subsampled cell populations, which are subject to network construction and form a multilayer scGRN. (**B**) *Network construction*. Principal component (PC) regression is used for scGRN construction; each scGRN is represented as a weighted adjacency matrix. (**C**) *Tensor denoising*. Two samples produce two multilayer GRNs, form two three-order tensors, which are subsequently decomposed into multiple components. Top components of tensor decomposition are then used to reconstruct two denoised multilayer scGRNs. Then, two denoised multilayer scGRNs are collapsed by taking average weight across layers, respectively. (**D**) *Manifold alignment*. The two single-layer average scGRNs are then aligned with respect to common genes using a nonlinear manifold alignment algorithm. Each gene is projected to a low-rank manifold space as two data points, one from each sample. (**E**) *Differential regulation test*. The distance between the two data points is the relative difference of the gene in its regulatory relationships in the two scGRNs. Ranked genes are subject to tests for their significance in differential regulation between scGRNs.

#### Numerical methods

The numerical methods used to construct and compare scGRNs involve the following five steps:

##### Step 1. Pre-processing data and subsampling cells

The input data are two scRNAseq expression data matrices, ***X*** and ***Y***, containing expression values for *n* genes in *m_1_* and *m_2_* cells from two different samples, respectively. Next, *m* cells in ***X*** and ***Y*** are randomly sampled to form ***X***′ and ***Y***′. This subsampling process is repeated *t* times to create two collections of subsampled cells 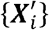 and 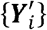, where *i* = 1, 2, …, *t*.

##### Step 2. Constructing initial networks

For each 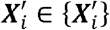, *i* = 1, 2, …, *t* principal component regression is used to construct a GRN. The constructed GRN from 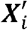 is stored as a weighted graph represented with an *n*×*n* weighted adjacency matrix 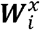. Similarly, for each 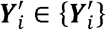, *i* = 1, 2, …, *t*, we construct a GRN using principal component regression and represent it with an *n*×*n* weighted adjacency matrix 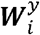. Diagonal values of each adjacency matrix are set to zeros, and other values are normalized by dividing by their maximal absolute value. Each normalized adjacency matrix is then filtered by retaining only the top 5% of edges ranked using the absolute egde weight, resulting in a sparse adjacency matrix.

##### Step 3. Denoising

Tensor decomposition [11] is used to denoise the adjacency matrices obtained in Step 2. The collection of *t* scGRNs for each sample, 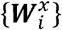 or 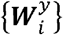, is processed separately as a third-order tensor, denoted as 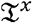 or 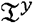, each containing *n*×*n*×*t* elements. The CANDECOMP/PARAFAC (CP) decomposition is applied to decompose 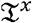 and 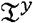 into components. Next, 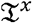 and 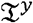 are reconstructed using top *r* components to obtain denoised tensors: 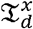 and 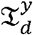. Denoised 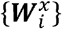 and 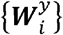 in 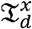 and 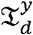 are collapsed by taking the average of edge weights for each edge to form two denoised, averaged matrices, 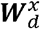 and 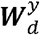, which are subsequently normalized as in step 2 and then symmetrized.

##### Step 4. Aligning genes onto a manifold

The weighted adjacency matrices 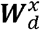 and 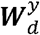 are regarded as two similarity matrices for a nonlinear manifold alignment procedure. The alignment is done by solving an eigenvalue problem with a Laplacian matrix derived from the joint matrices: 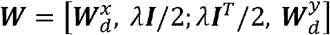, where *λ* is a tuning parameter and ***I*** is the identity matrix that reflects the binary correspondence between genes in the samples, ***X*** and ***Y***. As the result of manifold alignment, all genes in the samples, ***X*** and ***Y***, are projected on a shared, low dimensional manifold with a dimension *k_m_* ≪ *n*. The projections of each gene *j* from the samples, ***X*** and ***Y***, are two *k_m_*-dimensional vectors, 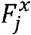 and 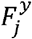.

##### Step 5. Ranking genes

For each gene *j*, let *d_j_* be the Euclidean distance between the gene’s two projections 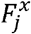 and 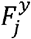 on the shared manifold: one is from the sample ***X***, and the other is from the sample ***Y***. Genes are sorted according to this distance. The greater the distance, the greater the regulatory shift.

In the following sections, we explain the rationale behind each step, the selection of machine learning methods, and implementation details.

#### Subsampling of cells

The rationale for randomly subsampling cells is close to that of ensemble learning, a technique where multi-model decisions are merged to improve overall performance. Instead of attempting to build a single scGRN, scTenifoldNet focuses on subsampling cells from a given scRNAseq expression matrix, building a number of ‘low-precision’ scGRNs from subsampled data sets, and then combining these scGRNs to obtain a high-precision scGRN. As mentioned above, current scRNAseq technology can produce the transcriptome profiles of thousands of cells from each sample. It is fundamentally difficult to process high-dimensionality and large-scale scRNAseq data, especially given that there can be substantial variation between cells even in a group of cells of the same type. For example, the presence of outlying cells, i.e., cells showing profiles of expression deviate from those of most other cells, can influence the construction of scGRNs. Therefore, subsampling offers promise as a technique for handling the noise in the input data sets. When the number of cells is small, the input data matrix may be resampled with replacement [12].

#### Constructing scGRNs using principal component regression

Although many GRN construction methods have been developed [1, 2, 4], it is unclear which one is suitable for constructing a large number of scGRNs from the subsampled data [9]. When dealing with multiple sets of input data, both the accuracy and computational efficiency of these algorithms have to be considered. We opted to use the PCNet method [5], which is based on principal component regression [13], after conducting a thorough review of the current methods. The principal component regression method extracts the first few (e.g., *k* = 3) principal components and then uses these components as the predictors in a linear regression model fitted using ordinary least squares. The values of the transformed coefficients of genes are treated as the strength and regulatory effect between genes to generate the network. The main use of principal component regression in scTenifoldNet lies in its ability to surpass the multicollinearity problem that arises when two or more explanatory variables are linearly correlated.

#### Denoising via low-rank tensor approximation

Removing the noise from constructed scGRNs is an important step of scTenifoldNet. Here the term noise is used in a broad sense to refer to any outlier or interference that is not the quantity of interest, i.e., the true regulatory relationship between genes. For each sample, the multilayer scGRN constructed from multiple subsampled data sets is regarded as a rank-three tensor. To reduce the noise in the multilayer scGRN, we decompose the tensor and reconstruct the multilayer scGRN using leading components. The idea is similar to that of denoising using truncated singular value decomposition (SVD). After cutting a larger portion of the noise spread over the lowest singular value components, the reconstructed data matrix based on the truncated SVD would, therefore, represent the original data with reduced noise. Indeed, tensor decomposition has been used in video data analyses for denoising and information extracting purposes [14]. It has also been used to impute missing data [15]. We use the CANDECOMP/PARAFAC (CP) algorithm [16] to factorize the two multilayer scGRNs separately and regenerate all adjacency matrices using leading components. The number of components used for reconstruction can be specified and is set to 3 by default. In the real data applications, we find the tensor GRN regeneration serves for two purposes: denoising and enhancing, i.e., making main signals stronger and making less important signals weaker.

#### Manifold alignment of two scGRNs

For a gene, its position in one of the two scGRNs (i.e., denoised adjacency matrices from the two samples) is determined by its regulatory relationships with all other genes. Here we regard each gene as a data point in a high-dimensional space where components of the data point are the features, i.e., weights between the gene and all other genes in the scGRN adjacency matrix. To compare the same gene’s positions in the two scGRNs, we first align the two scGRNs. To do so, we take a popular and effective approach for processing high-dimensional data, intuitively modeling the intrinsic geometry of the data as being sampled from a low-dimensional manifold—i.e., commonly referred to as the manifold assumption [17]. This assumption essentially means that local regions in the data can be mapped to low-dimensional coordinates, while the nonlinearity and high dimensionality in the data come from the curvature of the manifold. Manifold alignment produces projections between sets of data, given that the original data sets lie on a common manifold [18–21]. Manifold alignment matches the local and nonlinear structures among the data points from multiple sources and projects them to the same lowdimensional space while maintaining their local manifold structure of each source. The ability to flexibly learn and accurately represent the structure in the data with manifold alignment has been demonstrated in applications in automatic machine translation, face recognition, and so on [22, 23]. Here, we use manifold alignment to match genes in the two denoised scGRNs, one from each sample, to identify cross-network linkages. Consequently, the information of genes stored in two scGRNs is aligned, meaning points close together in the low-dimensional space are more similar than points that are farther apart.

#### Ranking genes and reporting DR genes

To identify genes whose regulatory status differs between the two samples, we calculate the distance between projected data points in the manifold alignment subspace. For each gene, if the gene appears in scGRNs of both samples, there are two data points for the same genes, one from each sample. We compute the Euclidean distance between the two data points of the gene and used the distance to measure the dissimilarity in the gene’s regulatory status in two scGRNs [24]. We do this for all genes shared between two samples and then rank genes by the distance. The larger the distance, the more different the gene in two samples. In this way, we obtain a list of ranked genes. These ranked genes are subject to functional annotation, such as using the pre-ranked Gene Set Enrichment Analysis (GSEA) [25] to assess the enriched functions associated with the top genes. To avoid choosing the number of selected genes arbitrarily, we compute *p*-values for genes using Chi-square tests, adjust *p*-values with a multiple testing correction, and select significant genes using the 10% FDR cutoff.

### Benchmarking the performance of scTenifoldNet using simulated data

#### Precision and recall of the network construction method adopted in scTenifoldNet

To show the effectiveness of principal component regression, we simulated scRNAseq data using a parametric method with a predefined scGRN model (see **Methods** for details). With these simulated data whose generating parameters are known, we compared the scGRNs constructed using principal component regression against the predefined scGRN (i.e., ground truth) to estimate the accuracy of principal component regression. Similarly, we tested the accuracy of other methods, including the Spearman correlation coefficient (SCC), mutual information (MI) [1], and GENIE3 (a random forest-based method [2, 7]). SCC and MI methods are computationally efficient, whereas GENIE3 is not, but GENIE3 is the top-performing method for network inference in the DREAM challenges [3]. For each method, their performances in recovering gene regulatory relationships were compared. Using the ground-truth interactions between genes generated according to pre-setting parameters, we found that the principal component regression method tended to produce more specific (better accuracy) and more sensitive (better recall) scGRNs than the other three methods across a wide range of settings of cell numbers in the input scRNAseq expression matrix (**Fig. 2A**). Principal component regression is also much faster than GENIE3. For instance, on a typical workstation, our implementation of principal component regression can construct a GRN for all-by-all 15,000 genes in less than 50 minutes, whereas GENIE3 requires more than 24 hours (data is available in **Supplementary Table S1**). This simulation study suggests that the principal component regression method is preferred to all other methods tested.

**Fig. 2.**
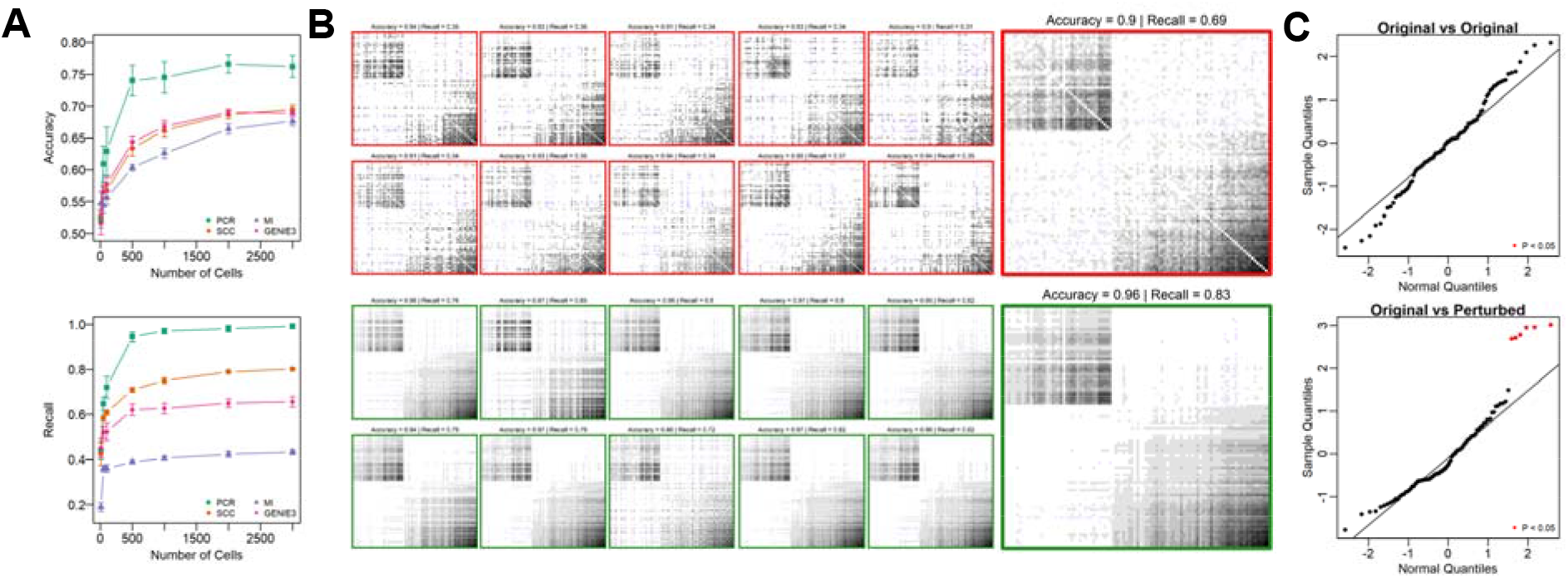
Benchmarking the performance of scTenifoldNet using simulated data. (**A**) The accuracy and recall of scGRN construction using different methods: principal component regression, SCC, MI, and GENIE3, as functions of the number of cells used in the analysis. Error bar is the standard deviation of the computed values after 10 bootstrapped evaluations. Accuracy is defined as = (TP+TN)/(TP+TN+FP+FN), and recall = TP/(TP+FN), where T, P, F, and N stands for true, positive, false and negative, respectively. PCR – principal component regression; SCC – Spearman correlation coefficient; MI – mutual information; GENIE3 – a random forest-based network construction method. (**B**) Visualization of the effect of tensor denoising on accuracy and recall of multilayer scGRNs. Each subpanel is a heatmap of a 100×100 adjacency matrix constructed using principal component regression over the counts of 500 randomly subsampled cells. Grayscale indicates the relative strength of regulatory relationships between genes. Top part includes networks before tensor denoising (adjacency matrices in heatmap with red box); bottom part includes corresponding networks after tensor denoising (adjacency matrices in heatmap with green box). In each part, adjacency matrices of networks of 10 subsamples (10 small heatmaps) and their average adjacency matrix (one large heatmap) are shown. (**C**) Evaluation of the sensitivity of scTenifoldNet in identifying punctual changes in the regulatory profiles. Top panel: evaluation of the original data matrix against itself; bottom panel: evaluation of the original matrix against the perturbed matrix. Significant genes identified using the differential regulation test (FDR < 0.05, the B–H correction) are indicated in red. All significant genes are perturbed in simulation and thus are expected to be identified.

#### Effect of denoising with tensor decomposition

To show the effect of tensor denoising, we simulated scRNAseq data (see **Methods**) and processed the data using the first two steps of scTenifoldNet, i.e., cell subsampling followed by the construction of scGRNs using principal component regression. We subsampled 500 cells each time and generated ten scGRNs. The ten scGRNs are treated as a multilayer network or a tensor to be denoised. For each scGRN, we kept the top 20% of the links. The presence and absence of links in each scGRN were compared with those in the simulated, ground-truth scGRN to estimate the accuracy of recovery and the rate of recall. **Fig. 2B** contains the heatmaps of adjacency matrices of the ten scGRNs before and after denoising (small panels). We also show two collapsed scGRNs (**Fig. 2B**, large panels), which were generated by averaging link weights across the ten scGRNs before and after denoising. These results illustrate the ability of scTenifoldNet to denoise multilayer scGRNs. For instance, tensor denoising improves the recall rate of regulatory relationships between genes by 25%. This simulation study suggests that tensor denoising could be useful for removing impacts of random dropout and other noise issues affecting the scGRN construction using scRNAseq data.

#### Detecting power illustrated with a simulated data set

We used simulated data to show the capability of scTenifoldNet in detecting differentially regulated (DR) genes. We first used the negative binomial distribution to generate a sparse synthetic scRNAseq data set (an expression matrix including 67% zeros in its values). This toy data set includes 2,000 cells and 100 genes. We called it sample 1. We then duplicated the expression matrix of sample 1 to make sample 2. We modified the expression matrix of sample 2 by swapping expression values of three randomly selected genes with those of another three randomly selected genes. Thus, the differences between samples 1 and 2 are restricted in these six genes. Using scTenifoldNet with the default parameter setting, we compared the originally generated expression matrix (sample 1) against itself (sample 1 vs. sample 1) and also against the manually perturbed version (sample 1 vs. sample 2). As expected, when comparing the original matrix against itself, none of the genes was identified to be significant. However, when samples 1 and 2 were compared, the six genes whose expression values were swapped were identified as significant DR genes (**Fig. 2C**, FDR < 0.05). These results are expected and support the sensitivity of scTenifoldNet in identifying subtly shifted gene expression programs.

### Real data analyses

#### Practical considerations of real data analysis using scTenifoldNet

First of all, we address several practical questions regarding the application of scTenifoldNet to real scRNAseq data. (1) What are the input expression matrices to be compared? The input to scTenifoldNet is two matrices of gene expression values (e.g., UMI counts) as measured in two samples to be compared. In each matrix, columns represent cells, and rows represent genes. We assume that each input matrix contains a sizable number of cells. For example, a typical input matrix may contain UMI counts for 5,000 genes and 2,000 cells. Whether a gene is expressed among cells can be determined by examining if this gene has a nonzero UMI count in more than 5% of cells. (2) How does scTenifoldNet handle cell heterogeneity? Heterogeneity in expression among cells is inevitable. scTenifoldNet is designed to tolerate a certain level of such heterogeneity as long as cells are of the same type. scTenifoldNet is not a data preparation tool. It also does not perform any clustering analysis for cells; it does not assign cells into cell types. We assume all cells in both input matrices are of the same type. Otherwise, the results would be difficult to interpret. (3) What if the number of cells is too small? We expect that each input matrix contains a sizable number of cells (e.g., n>2,000). If this is the case, the jackknife method (subsampling without replacement) is adapted by default: *m*=500 cells are subsampled each time. Alternatively, an *m*-out-of-*n* bootstrap method (subsampling with replacement) can be used [12]. When the number of cells is small (e.g., *n*=500), a full bootstrap method can be used, i.e., resampling 500 cells each time out of 500 given cells with replacement [12, 26]. (4) What is the relationship between scTenifoldNet analysis and DE analysis? scTenifoldNet analysis should be used as a complementary analysis method in addition to DE analysis, rather than replacing DE analysis. DE analysis is still the most widely used method for understanding the difference between two scRNAseq samples. scTenifoldNet is designed based on a different principle from that is underlying DE analysis. Thus, the results of scTenifoldNet analysis and DE analysis are not supposed to be compared side by side. It is not uncommon that scTenifoldNet and DE analyses report same genes to be significant. This is because the change of the regulatory pattern of a gene in scGRNs may be associated with the change of the gene’s expression level.

#### Analysis of transcriptional responses of neurons to acute morphine treatment

To illustrate the use of scTenifoldNet, we first applied scTenifoldNet to a scRNAseq data set from [27]. This is a study on transcriptional responses of mouse neural cells to morphine (**Fig. 3A**). In the study [27], Avey and colleagues performed scRNAseq experiments with the nucleus accumbens (NAc) of mice after four hours of the morphine treatment, using mice treated with saline as mock controls. Single-cell expression data was obtained for 11,171 and 12,105 cells from four morphine- and four mock-treated mice, respectively [27]. The measured cells were clustered to identify neurons (7,972 and 8,912 from morphine- and mock-treated samples, respectively); the identified neurons were then sub-grouped into 11 clusters, including major clusters of D1 and D2 medium spiny neurons (MSNs), comprising ~95% of the neurons in the NAc. Using differential expression (DE) analysis implemented in SCDE [28], Avey et al. identified several hundred genes that are differentially expressed between morphine- and mock-treated samples (**Supplementary Table S2** of [27]). Although this result is intriguing, we argue that it seems that when so many genes are identified as ‘significant players,’ it is difficult to interpret the result and to pinpoint the specific regulatory mechanism underlying the true response. Indeed, instead of performing functional enrichment analysis with identified DE genes, the subsequent analyses in the study of [27] were re-focused on a tiny portion of D1 MSNs, called activated MSNs. It is only when activated MSNs were compared to all other D1 MSNs that 256 DE genes were identified (SCDE, P < 0.001, **Supplementary Table S2** of [27]). These genes were then found to be associated with several terms related to opioid addiction, including morphine dependence and opioid-related disorders (**Supplementary Table S3** of [27]). In the morphine-treated sample, less than 4.5% of D1 MSNs are activated MSNs; in the mock-treated sample, less than 2% (see **Fig. 2B** of [27]). In view of these, we point out here that while relevant signals can be detected using traditional DE analysis, the analytical method involves extensive human intervention—i.e., an iterative clustering procedure is needed to identify a final population of cells (in this case, activated MSNs). The cell population size is small, making the analysis result potentially variable.

**Fig. 3.**
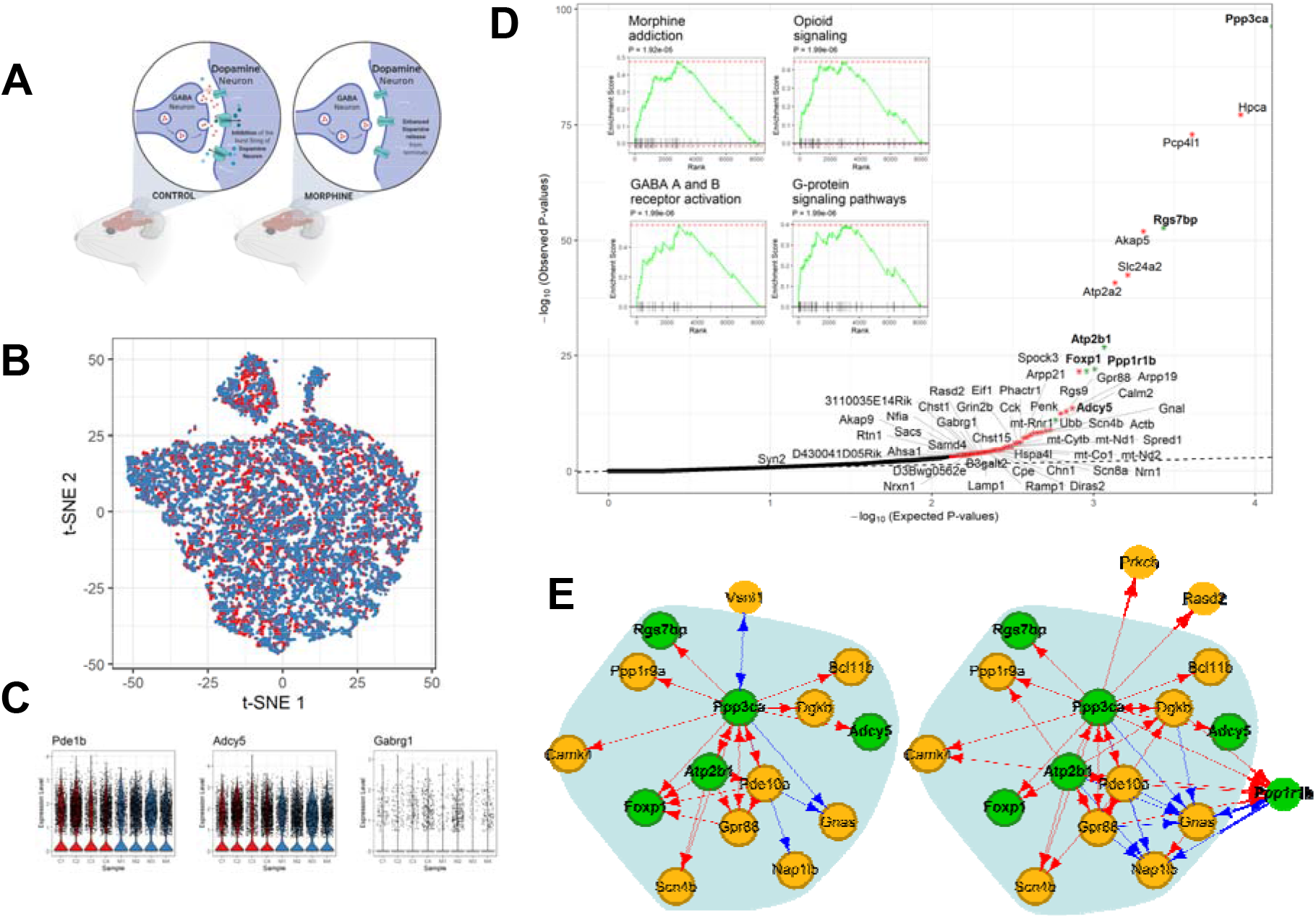
Analysis of transcriptional responses to morphine in mouse cortical neurons. (**A**) Illustration of experimental design and data collection of the morphine response study [27]. (**B**) A t-SNE plot of 7,972 and 8,912 neurons from morphine-treated (blue) and mock-treated (red) mice, respectively. (**C**) Violin plots show the log-normalized expression levels of representative DR and/or DE genes in four (M) morphine- and four (C) mock-treated mice. (**D**) Quantile-quantile (Q-Q) plot for observed and expected p-values of the 8,138 genes tested. Genes (n=65) with FDR < 0.1 are shown in red; genes (n=56) with FDR < 0.05 are labeled with asterisk. Inset shows results of the GSEA analysis for genes ranked by their distances in manifold aligned scGRNs from morphine- and mock-treated mice. (**E**) The module enriched with DR genes and the corresponding subnetworks in two scGRNs. For illustrative purposes, the module is centered on the DR gene, *Ppp3ca*. Significant DR genes (FDR<0.05) in the module are highlighted in green. Edges are color-coded: red indicates a positive association, and blue indicates negative. Weak edges are filtered out by thresholding for clear visualization, and the background shadow indicates the shared portion of the module in the two scGRNs.

We were motivated by these considerations and set out to re-analyze the data. We first reproduce the results of the DE analysis. We found that the mock- and morphine-treated neurons indeed exhibited a striking similarity. For example, mock- and morphine-treated neurons are indistinguishable in a tSNE plot (**Fig. 3B**); expression levels of several known morphine responsive genes, e.g., *Adcy5, Ppp1r1b*, and *Ppp3ca*, show no difference (**Fig. 3C**). Thus, a direct comparison of gene expression between neurons using the DE method may have limited power to identify relevant genes involved in the morphine response.

Next, using scTenifoldNet, we identified 56 genes showing significant differences in their transcriptional regulation between mock- and morphine-treated neurons, indicated by greater distances between genes’ positions in the aligned manifold of two scGRNs (FDR < 0.05, Chisquare test with B-H multiple test adjustment, see **Methods** for details). These genes are: ***Ppp3ca**, **Hpca**, Pcp4l1, Rgs7bp, **Akap5**, Slc24a2, Atp2a2, **Atp2b1**, **Ppp1r1b**, Foxp1, Spock3, Arpp19, **Gpr88**, **Rgs9**, **Adcy5**, **Gnal**, Ubb, **Scn4b**, Actb, **Calm2**, **Penk**, mt-Rnr1, **Arpp21**, Phactr1, Cck, **Eif1**, mt-Nd1, mt-Cytb, Spred1, mt-Nd2, mt-Co1, Hspa4l, Nrn1, **Scn8a**, **Chn1**, Diras2, Cpe, **Ramp1**, B3galt2, **Chst15**, **Grin2b**, Lamp1, **Rasd2**, Gabrg1, Chst1, 3110035E14Rik, Nfia, Samd4, Sacs, Nrxn1, **D3Bwg0562e**, Akap9, Rtn1, Ahsa1, D430041D05Rik*, and *Syn2* (genes sorted according to the significance level, see also **Fig. 3D**). The pre-ranked GSEA analysis [25] showed that these genes are enriched for *opioid signaling, signaling by G protein-coupled receptors, reduction of cytosolic Calcium levels*, and *morphine addiction* (**Fig. 3D** inserts). It is known that morphine binds to the opioid receptors on the neuronal membrane. The signal is then transmitted through the G-protein signaling system, inhibiting the adenylyl cyclase in the cytoplasm and decreasing the levels of cAMP and the calcium-channel conduction [29–31]. Furthermore, 21 (highlighted in bold) of the 56 identified DR genes were found to be targets of RARB (38%, adjusted P-value < 0.01, enrichment test by Enrichr [32, 33] based on results of ChIP-seq studies [34]). RARB encodes for a plastic TF playing a role in synaptic transmission in dopaminergic neurons and the adenylate cyclase-activating dopamine receptor signaling pathway [35, 36]. Thus, these enriched functions are relevant to the morphine stimulus, which is known to induce the disinhibition of dopaminergic neurons by GABA transmission, enhance dopamine release, and cause addiction [37, 38]. Using the constructed scGRN, we were able to trace DR genes back to their topological positions in the network and examine their interacting genes. **Fig. 3E** shows such a network module, including multiple DR genes.

In this case, scTenifoldNet is used as an unsupervised tool, and no human interference is needed to operate. This feature is critical when referring to this specific set of data because where the signal is limited to rare types of cells, there is a chance that a less sensitive approach would miss the signal, especially when human interference is not provided. It is ideal to have an unsupervised tool that is sensitive to signals, and robust to variation between cells at the same time. We note that scTenifoldNet is a different tool to conventional DE analysis tool—scTenifoldNet reported less DR genes in terms of the number of genes, compared with DE genes identified in the original study [27]. Among the 56 DR genes that scTenifoldNet detected, 11 (*Actb*, *Adcy5, Akap9, D430041D05Rik, Eif1, Pcp4l1, Penk, Phactr1, Rasd2, Scn4b* and *Ubb*) are among the 256 DE genes reported in [27](see **Supplementary Table S2** of [27]). The number of overlap genes is not significantly higher than expected by random according to a hypergeometric test (*P* = 0.29) with a total of 1,432 genes (from **Supplementary Table S2** of [27]) included in the test. **Fig. 3C** shows expression levels of three representative genes: *Pde1b, Adcy5* and *Gabrg1*, in neurons from mock- and morphine-treated mice. All three genes are known to be involved in morphine response [39–41], but only when DE and DR tests are applied jointly, all three genes are identified: *Pde1b* is a DE but not a DR gene, *Adcy5* a DR and DE gene, and *Gabrg1* a DR but not DE gene.

#### Analysis of transcriptional responses of a carcinoma cell line to cetuximab

To further illustrate the power of scTenifoldNet in identifying genes associated with specific perturbations, we applied scTenifoldNet to another published scRNAseq data [42]. In this study, Kagohara *et al*. [42] use scRNAseq to study mechanisms that lead the development of resistance to cetuximab in head and neck squamous cell carcinoma (HNSCC)(**Fig. 4A**). Cetuximab is a human-murine chimeric monoclonal antibody used for the treatment of metastatic colorectal cancer, metastatic non-small cell lung cancer, and head and neck cancer. In conjunction with the radiotherapy, cetuximab improves the objective response rate in first-line treatment of recurrent or metastatic squamous cell carcinoma of the head and neck [43]. Cetuximab binds to the extracellular domain of the epidermal growth factor receptor (EGFR) on both normal and tumor cells [44]. EGFR is over-expressed in many cancers. Competitive binding of cetuximab to EGFR blocks the phosphorylation and activation of receptor-associated kinases and their downstream targets, e.g., MAPK, PI3K/Akt, and Jak/Stat pathways [45], thereby reducing their effects on cell growth and metastatic spread. It is known that blocking EGFR activation also affects cellular processes such as apoptosis, cell growth, and vascular endothelial growth factor (VEGF) production [46]. Cetuximab is also known to cause degradation of the antibody-receptor complex and the downregulation of *EGFR1* expression [47].

**Fig. 4.**
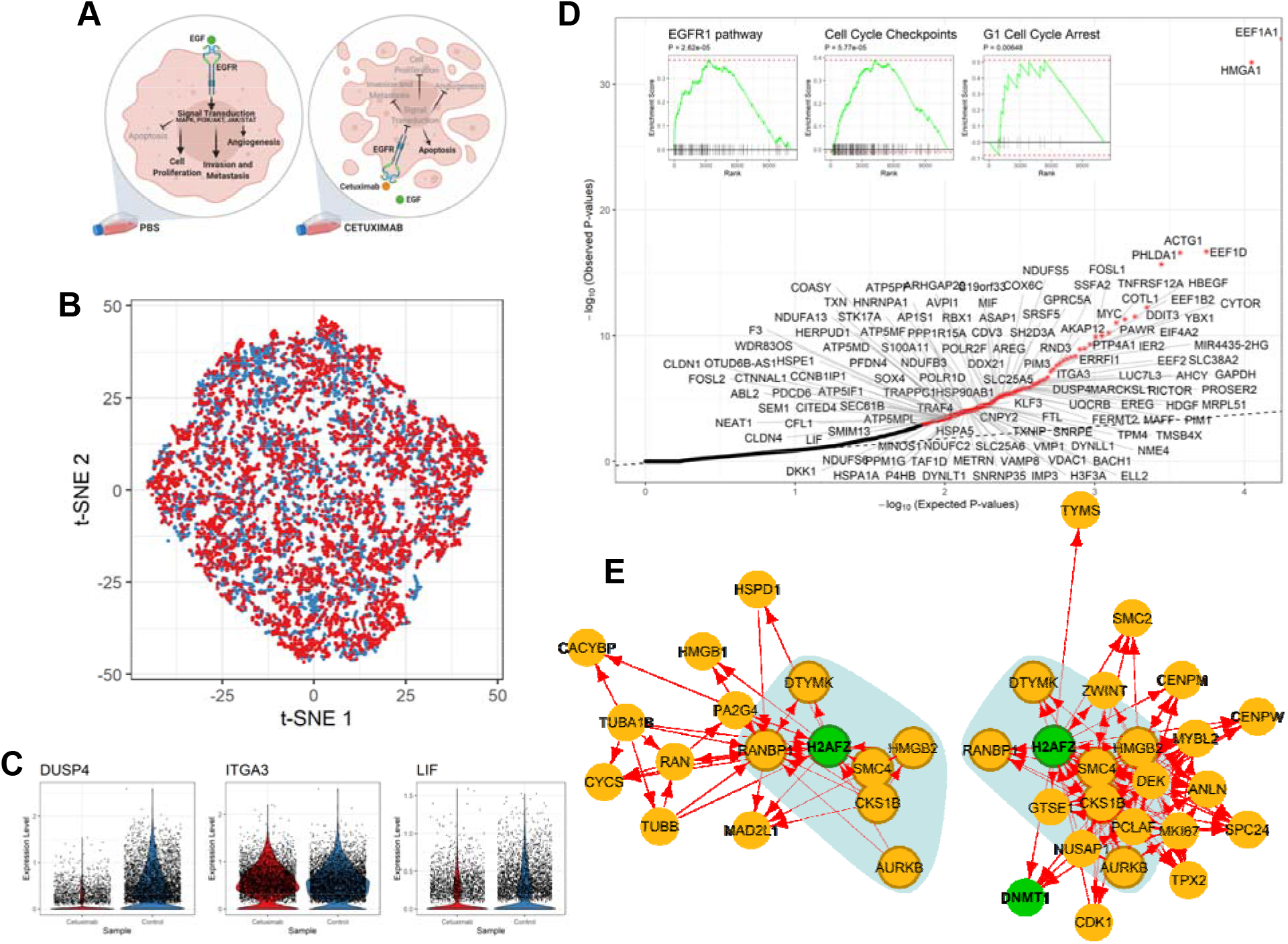
Analysis of transcriptional responses of a carcinoma cell line to cetuximab. (**A**) Illustration of experimental design, including sample groups and the known mechanism of drug action, in the study of cetuximab resistance of HNSCC cell lines [42]. (**B**) t-SNE plot of 5,217 and 4,507 HNSCC-SCC6 cells treated with cetuximab (red) and PBS (blue), respectively. (**C**) Violin plots show the log-normalized expression levels of selected DR genes in SCC6 cells with and without cetuximab treatment. (**D**) Q-Q plot for observed and expected p-values of the 7,503 genes tested. Genes (*n* = 25) with FDR < 0.05 are labeled with asterisk. Inset shows the results of the GSEA analysis for genes ranked by their distances in manifold aligned scGRNs from young and old mice. (**E**) A representative module with DR genes and corresponding subnetworks in two scGRNs. The module is enriched with DR genes and the corresponding subnetworks in two scGRNs. For illustrative purposes, the module is centered on the DR gene, *H2AFZ*. The colors, edges, and marks are presented as in **Fig. 3E**.

Kagohara *et al*. [42] sequenced the transcriptome profile of cells before and after Cetuximab treatment for 120 hours in three different HNSCC cell lines: SCC1, SCC6, SCC25. They found that SCC6 is the most sensitive to the cetuximab treatment, reporting 8,389 genes as differentially expressed (including 4,166 upregulated and 4,223 downregulated ones with P < 0.05; **Supplementary Table S4** of [42]). Such a large number of differentially expressed genes makes it difficult to identify genes directly associated with the molecular mechanism through which cetuximab acts.

We extracted scRNAseq data for 4,507 and 5,217 SCC6 cells treated with and without cetuximab, respectively (**Fig. 4B**). Expression levels of three genes, *DuSP4, TIGA3* and *LIF*, in cells of two treatment groups, are shown in **Fig. 4C**. All three genes are in the EGFR pathway. We used scTenifoldNet to re-analyze the data and identified 125 differentially regulated genes (FDR<0.05, **Fig. 4D**). These genes are: *EEF1A1, HMGA1, EEF1D, ACTG1, PHLDA1, **HBEGF**, EEF1B2, **COTL1**, **TNFRSF12A**, DDIT3, **MYC**, **FOSL1**, **PAWR**, YBX1, CYTOR, **EIF4A2**, **PTP4A1**, IER2, MIR4435-2HG, AKAP12, SSFA2, **ERRFI1**, EEF2, **ITGA3**, **SLC38A2**, LUC7L3, AHCY, GAPDH, PROSER2, RICTOR, MARCKSL1, DUSP4, MRPL51, **HDGF**, **EREG**, **PIM1**, RND3, PIM3, **GPRC5A**, UQCRB, MAFF, FERMT2, SRSF5, **SH2D3A**, NDUFS5, SLC25A5, COX6C, AREG, ASAP1, MIF, C19orf33, **CDV3**, DDX21, RBX1, **AVPI1**, POLR2F, **KLF3**, TMSB4X, FTL, **TPM4**, NME4, **SNRPE**, ARHGAP29, **PPP1R15A**, AP1S1, ATP5PF, **HSP90AB1**, **POLR1D**, DYNLL1, CNPY2, **BACH1**, TXNIP, **ELL2**, VMP1, VDAC1, H3F3A, IMP3, VAMP8, SLC25A6, SNRNP35, METRN, DYNLT1, **HSPA5**, HNRNPA1, NDUFB3, S100A11, ATP5MF, NDUFC2, **STK17A**, **TXN**, COASY, TAF1D, NDUFA13, P4HB, PPM1G, HSPA1A, HERPUD1, ATP5MD, PFDN4, SOX4, TRAPPC1, TRAF4, **F3**, WDR83OS, **HSPE1**, CCNB1IP1, MINOS1, OTUD6B-AS1, ATP5IF1, **CTNNAL1**, **CLDN1**, **PDCD6**, **FOSL2**, **SEC61B**, **ABL2**, CITED4, SEM1, ATP5MPL, NEAT1, CFL1, NDUFS6, **CLDN4**, SMIM13, **LIF***, and *DKK1*.

This gene list is enriched with genes (39/125, names shown in bold) that are under the regulation of TFs: *SMAD2* and *SMAD3*. The pre-ranked GSEA analysis [25] shows that these DR genes are associated with *EGFR1* pathway (*DUSP4, ITGA3, LIF, DDX21, AREG, EREG, CLDN4, MYC, COTL1, TXNIP, LUC7L3, PHLDA1, HBEGF), regulation of apoptosis (HSP90AB1, HSPA5, MIF, YBX1, EEF2, DYNLL1, COX6C, HSPE1, MYC, DDIT3, PIM1, VDAC1, SEC61B, SLC25A5, PFDN4, HNRNPA1, GAPDH, IER2, HSPA1A, EEF1A1, DUSP4, MAFF, CITED4, CLDN1, PHLDA1, AREG, RBX1), cell cycle checkpoints*, and *G1 cell cycle arrest* (**Fig. 4D**, inserts; due to limited space, GSEA result of *regulation of apoptosis* is not shown). Once again, scTenifoldNet identified a much smaller set of significant genes compared to those reported in the original paper [42]: 125 DR genes vs. 8,389 DE genes. Nevertheless, functional analyses show that scTenifoldNet identified a more specific gene set relevant to cetuximab’s mechanism of action. Further scrutinization of enriched molecular functions of these DR genes will help to identify more regulatory targets induced by cetuximab in HNSCC cells.

#### Analysis of transcriptional responses of alveolar type 1 cells to *Nkx2-1* gene knockout

In the third example, we applied scTenifoldNet to another published scRNAseq data from type 1 alveolar (AT1) cells [48]. AT1 cells are responsible for gas exchange, the physiological function of the lung [49]. Little et al., [48] found that NK homeobox 2-1 (*Nkx2-1*) is expressed in AT1 cells and thought *Nkx2-1* might be essential to the development and maintenance of AT1 cells. To determine the function of NKX2-1 during the development of AT1 cells, they performed scRNAseq experiment to obtain the transcriptome profile of cells from lungs of *Nkx2-1^CKO/CKO^; Aqp5^Cre/+^* mutant mice (i.e., the knockout [KO] mice) and littermate controls (i.e., the wild-type [WT] mice). They used early infant mice (postnatal day 10, P10) because P10 represents an intermediate time point when individual AT1 cells in the mutant lung are expected to collectively feature the full range of transcriptomic phenotypes. They reported 3,622 DE genes (2,105 upregulated and 1,517 downregulated, **Supplementary Dataset S1** of [48]) between the KO and WT mice. Their analyses suggest that, without *Nkx2-1*, developing AT1 cells lose their molecular markers, morphology, and cellular quiescence, leading to aberrant expression of gastrointestinal (GI) genes, alveolar simplification and lethality (**Fig. 5A**).

**Fig. 5.**
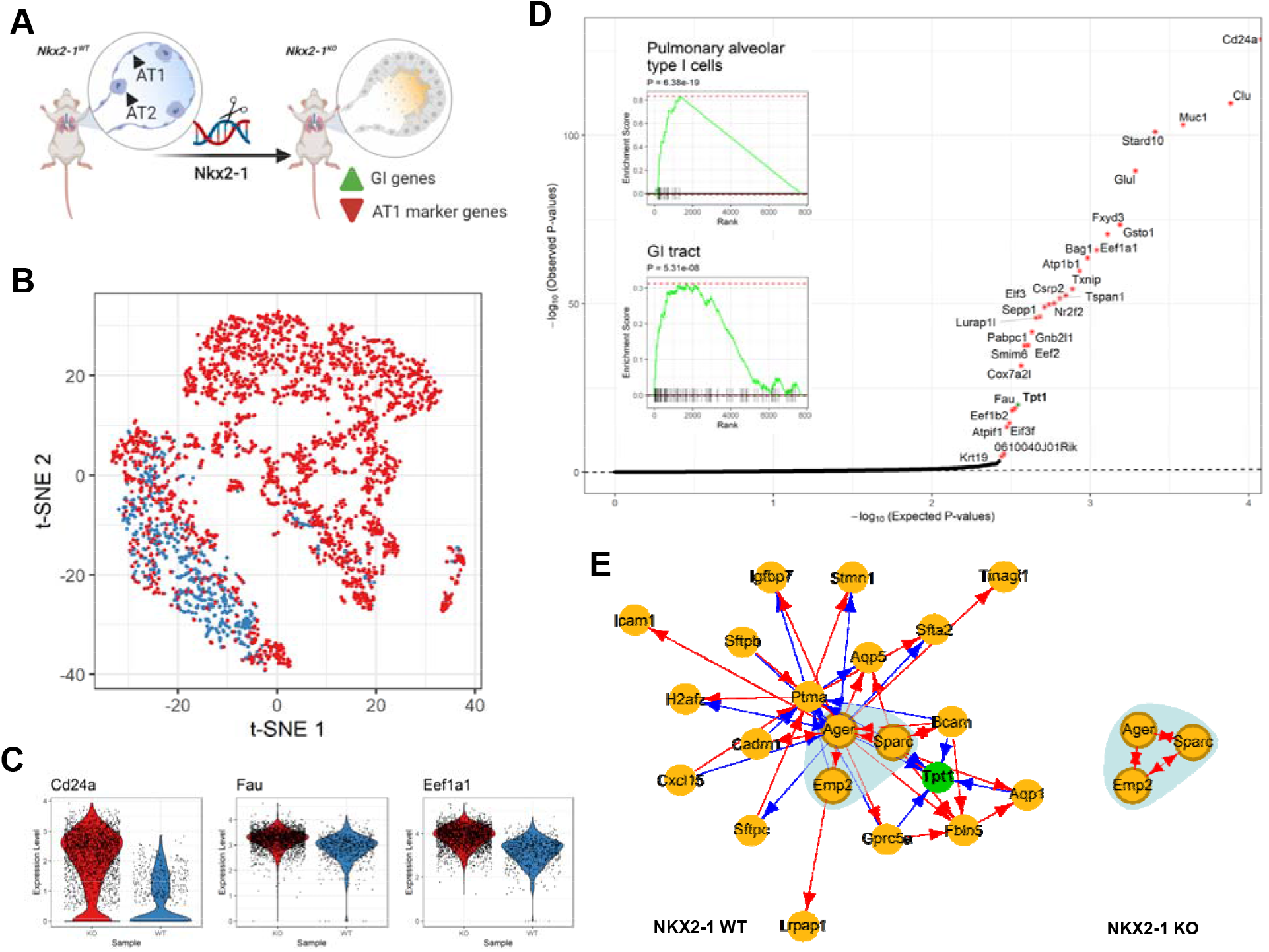
Analysis of transcriptional responses of alveolar type 1 cells to Nkx2-1 gene knockout. (**A**) Illustration of experimental design and data collection of the KO experiment [48]. (**B**) t-SNE plot of 2,397 and 638 AT1 cells from Nkx2-1 KO mice (red) and WT mice (blue). (**C**) Violin plots show the log-normalized expression levels of selected DR genes in KO (red) and WT (blue) mice. (**D**) Q-Q plot for observed and expected p-values of tested genes. Genes (*n*=29) with FDR < 0.05 are labeled with asterisk. Inset shows the results of the GSEA analysis for genes ranked by their distances in manifold aligned scGRNs. (**E**) A representative module that contains DR gene, *Tpt1*, in the WT mice. Most parts of the module disappear in the KO mice. The colors, edges and marks are presented as in **Fig. 3E**.

To evaluate the power of scTenifoldNet in identifying regulatory changes caused by gene KO, we re-analyzed the transcriptional profiles of 2,397 mutant AT1 cells from the *Nkx2-1^CKO/CKO^; Aqp5^Cre/+^* mice and 638 AT1 cells from the WT mice (**Fig. 5B**). Expression levels of *Cd24a, Fau* and *Eef1a1* in AT1 cells of KO and WT mice are shown in **Fig. 5C**. *Cd24a* is a marker gene for AT1 cells; *Fau* and *Eef1a1* are GI genes, known to be highly expressed in the GI tissues. Using scTenifoldNet, we identified 29 genes exhibiting significant difference in their regulation between the two samples: KO vs. WT (FDR < 0.05; ***Cd24a**, Clu, **Muc1**, Stard10, Glul, **Fxyd3**, **Gsto1**, Eef1a1, Bag1, **Atp1b1**, Txnip, Csrp2, **Tspan1**, Nr2f2, **Elf3**, Sepp1, Pabpc1, Lurap1l, Gnb2l1, Eef2, Smim6, Cox7a2l, **Tpt1**, Fau, Eef1b2, Eif3f, Atpif1, **0610040J01Rik, Krt19***)(**Fig. 5D**). This gene list is enriched with genes under regulation of TF, *Sox2* (FDR < 0.05; labeled in bold) [50]. The gene list is also enriched with genes highly expressed in the intestine (FDR < 0.05; *Gsto1, Stard10, Atpif1, Atp1b1, Eef2, Clu, Gnb2l1, Eef1b2, Eef1a1, Muc1, Cox7a2l, Krt19, Elf3, Fxyd3, Bag1, Txnip, Pabpc1, Fau, Eif3f, Glul, Tspan1, Tpt1*) and the gut (FDR < 0.05; *Gsto1, Stard10, Atpif1, Nr2f2, Atp1b1, Eef2, Clu, Gnb2l1, Eef1b2, Eef1a1, Muc1, Cox7a2l, Krt19, Csrp2, Lurap1l, Fxyd3, Pabpc1, Fau, Eif3f, Glul, Tspan1, Tpt1*), which is in agreement with the reported by the authors of the data set [48]. Using pre-ranked GSEA analysis [25], we were able to detect the effect of *Nkx2-1* KO on the cellular identity of AT1 cells, shown by the significant enrichment of gastrointestinal marker genes (*Ager, Col4a4, Col4a3, Aqp5, Emp2, Crlf1, Icam1, Egfl6, Vegfa, Gprc5a, Pdpn, Cldn18, Scnn1g, Akap5, Hopx, Scnn1b, Scnn1a, Sec14l3, Sema3e, Krt7, Gramd2, Clic5, Mmp11, Ctgf, Rtkn2, Pxdc1, Sema3b, Fstl3*) [51] (**Fig. 5D**, inserts).

#### Analysis of transcriptional responses of human dermal fibroblasts to the doublestranded RNA stimulus

Next, we show the use of scTenifoldNet to a scRNAseq data set from human dermal fibroblasts [52]. In the original paper, Hagai et. al. [52] focused on single-cell transcriptional responses induced by the stimulus of polyinosinic-polycytidylic acid (polyI:C), a synthetic double-stranded RNA (dsRNA)(**Fig. 6A**). They obtained and compared transcriptomes of 2,553 unstimulated and 2,130 stimulated cells and identified 875 DE genes (**Table S3** of [52]). These DE genes include *IFNB, TNF, IL1A*, and *CCL5*, encoding antiviral and inflammatory gene products, and are enriched for *inflammatory response, positive regulation of immune system process*, and *response to cytokine*, among many others biological processes and pathways. We found the original scRNAseq data has a batch effect between two samples, but the global batch effect can be removed using Harmony [53], as shown in the tSNE-plot of cells of two samples (**Fig. 6B**). Nevertheless, the differences in the expression level between samples can still be detected in selected genes with Harmony-processed data (**Fig. 6B**).

**Fig. 6.**
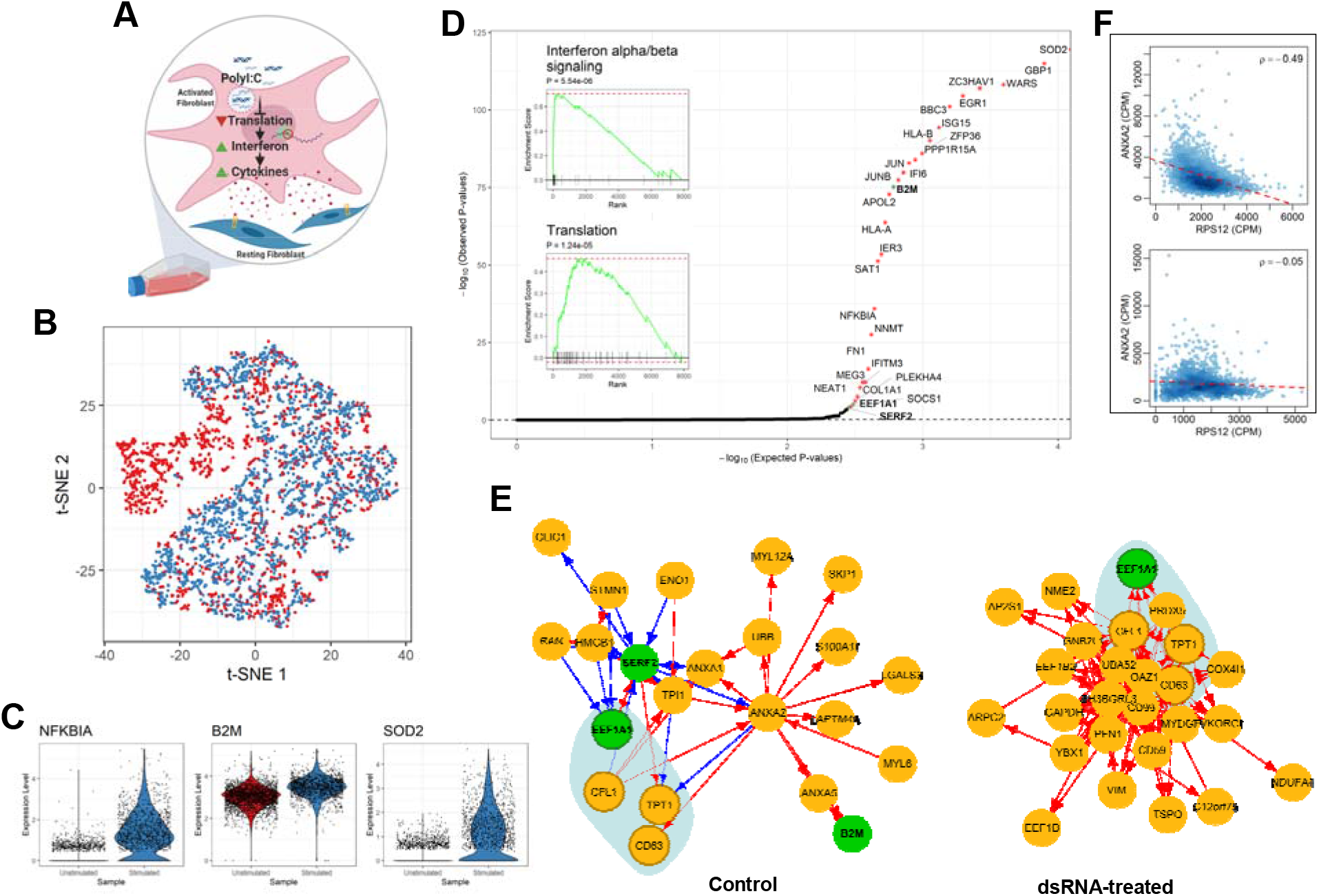
Analysis of transcriptional responses of human dermal fibroblasts to the double-stranded RNA stimulus. (**A**) Illustration of experimental design and tested mechanism of transcriptional responses [52]. (**B**) t-SNE plot of human dermal fibroblasts before (blue) and after (red) dsRNA stimulus. (**C**) Violin plots show the log-normalized expression levels of selected DR genes before (blue) and after (red) stimulus. (**D**) Q-Q plot for observed and expected p-values of tested genes. Genes (*n* = 29) with FDR < 0.05 are labeled with asterisk. Inset shows the results of GSEA analysis for genes ranked by their distances in manifold aligned scGRNs. (**E**) Comparison of a representative module that contains three DR genes in the control sample. The colors, edges and marks are presented as in **Fig. 3E**. (**F**) Scatter plots show the correlation between *TPT1* and *ANXA2* before (top) and after (bottom) dsRNA stimulus.

Applying scTenifoldNet to the processed data, we identified 29 DR genes: ***SOD2**, **GBP1**, WARS, ZC3HAV1, **EGR1**, **BBC3**, ISG15, **HLA-B**, ZFP36, **PPP1R15A**, **JUN**, IFI6, **JUNB**, **B2M**, APOL2, **HLA-A**, **IER3**, SAT1, **NFKBIA**, **NNMT**, **FN1**, IFITM3, MEG3, NEAT1, COL1A1, PLEKHA4, EEF1A1, SOCS1, and SERF2* (**Fig. 6D**). Among them, 14 (highlighted in bold) are targets of TF, *RELA* [54] (48%, adjusted *p*-value < 0.01, enrichment test by Enrichr [32, 33]). These DR genes are functionally enriched for *interferon signaling (IFITM3, EGR1, SOCS1, IFI6, HLA-B, ISG15, HLA-A, GBP1, B2M), immune system (IFITM3, NFKBIA, EGR1, JUN, SOCS1, IFI6, HLA-B, ISG15, HLA-A, GBP1, B2M), interleukin-1 regulation of extracellular matrix (NFKBIA, ZFP36, JUN, HLA-B, SOD2*), and among others.

Once again, scTenifoldNet reports fewer genes than DE analysis does in the original paper [52]. Through comparing DR genes with the DE genes, we found that enriched functions of DE genes reflect a final status of cells after cells responding to the dsRNA stimulus, whereas the enriched functions of DR genes reflect the activities associated with ongoing regulatory processes and immune responses to the stimulus. In this sense, DR genes are valuable for informing of mechanisms, through which the dsRNA acts to induce immunological responses [55–57]. For example, it is known that the dsRNA inhibits the translation of mRNA to proteins [56] and leads the synthesis of interferon, which induces the synthesis of ribosomal units that are able to distinguish between cell mRNA and viral RNA [57]. Interferon also promotes cytokine production that activates the immune responses and induces inflammation [55]. To further illustrate the changes in the regulatory patterns between samples, we plotted the GRN module around the EEF1A1 gene. It can be seen that, before and after the dsRNA treatment, the interacting partnership of the genes is changed substantially (**Fig. 6E**). Two scatter plots show the change of correlation between *TPT1* and *ANXA2*, as an example (**Fig. 6F**). The negative correlation between the two genes’ expression among cells disappears after the dsRNA treatment and, thus, the two genes are not linked in the scGRN constructed using the after-treatment data.

#### Analysis of transcriptional responses of mouse neurons in Alzheimer’s disease

Lastly, we applied scTenifoldNet to scRNAseq data of isolated single nuclei from the brains of the WT and 5xFAD mice [58]. The 5xFAD strain recapitulates the major features of Alzheimer’s disease amyloid pathology. The genotype of these mice contains several Familial Alzheimer’s Disease (FAD) mutations in *APP* and *PSEN1*, causing the overexpression of mutant human amyloid-beta (Aβ) precursor protein and human presenilin 1. The 5xFAD model rapidly develops amyloid pathology, with high levels of intraneuronal Aβ accumulation beginning around 1.5 months of age, and extracellular Aβ deposition beginning around two months [59].

In the original paper [58], Zhou et al. compared single-cell gene expression between 6-month-old WT mice with 6-month-old_5xFAD mice. They found that neurons show limited responses to Aβ peptides—compared to microglia and oligodendrocytes, neurons show minimal transcriptional changes (149 DE genes) between WT and 5xFAD mice. To test whether scTenifoldNet can detect genes whose expression is differentially regulated between WT and 5xFAD mice, we decided to apply our method to this scRNAseq data, exclusively in neurons. We downloaded expression data matrices from the GEO database using accession number GSE140511 and extracted expression data of neurons from two samples: GSM4173505 (WT2) and GSM4173511 (WT_5XFAD2)(**Fig. 7A**).

**Fig. 7.**
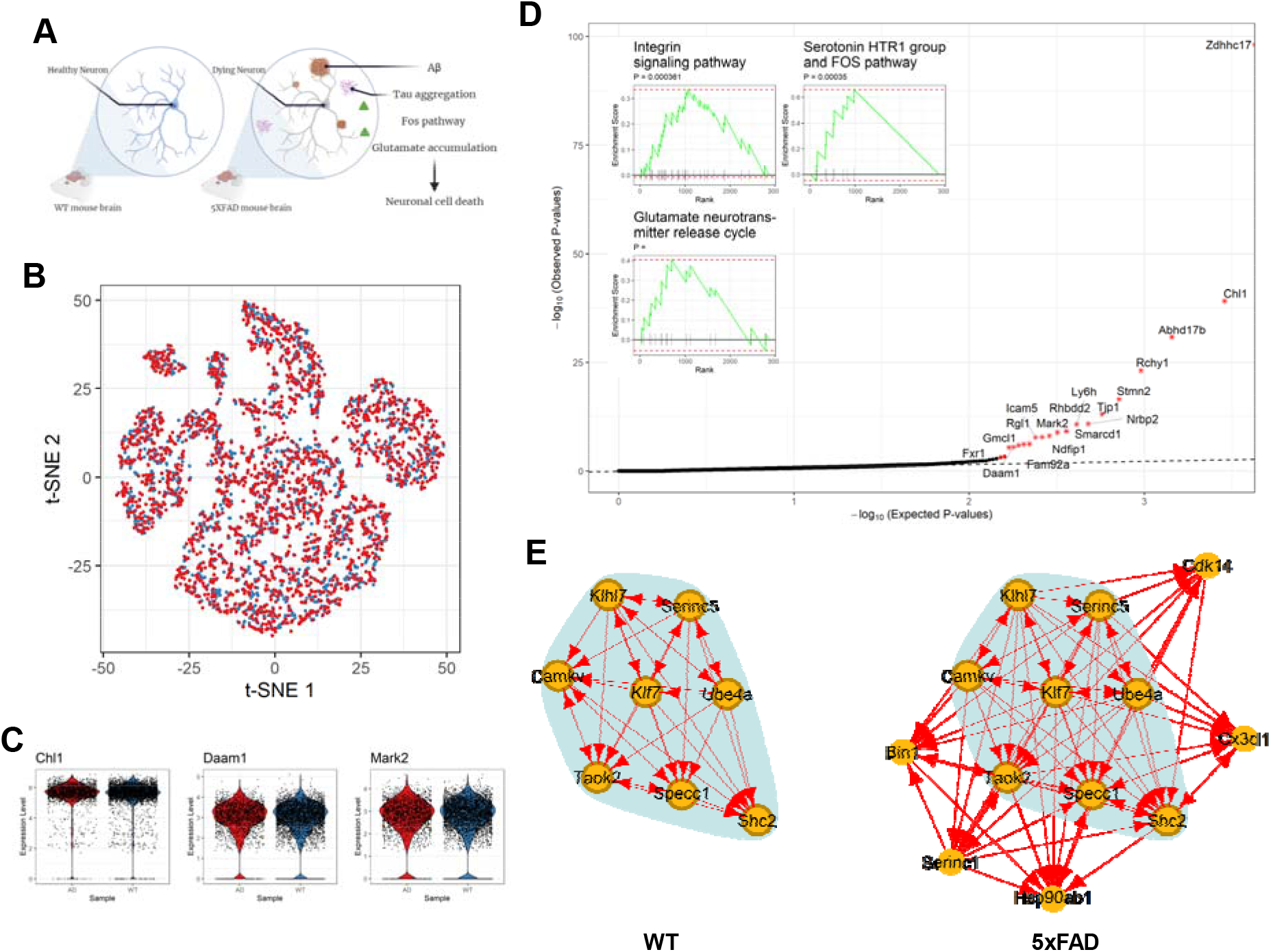
Analysis of transcriptional responses of neurons to amyloid-beta (Aβ) peptides in the 5xFAD mice, a model of Alzheimer’s disease. (**A**) Illustration of experimental design and data collection of the 5xFAD mice study [58]. (**B**) t-SNE plot of neurons of the 5xFAD (red) and WT (blue) mice. (**C**) Violin plots show the log-normalized expression levels of selected DR genes in neurons of the 5xFAD (red) and WT (blue) mice. (**D**) Q-Q plot for observed and expected p-values of tested genes. Genes (*n*=18) with FDR < 0.05 are labeled with asterisk. Inset shows the results of the GSEA analysis for genes ranked by their distances in manifold aligned scGRNs. (**E**) Comparison of a representative module that contains topranked DR genes between the two scGRNs. The colors, edges and marks are presented as in **Fig. 3E**.

After re-analyzing the data using scTenifoldNet, we identified 18 DR genes: *Zdhhc17, Chl1, Abhd17b, Rchy1, Stmn2, Tjp1, Nrbp2, Ly6h, Smarcd1, Rhbdd2, Ndfip1, Mark2, Icam5, Fam92a, Rgl1, Gmcl1, Daam1*, and *Fxr1* (sorted by significance, FDR<0.05, **Fig. 7D**). For functional enrichment analysis, we relaxed the significant-gene cutoff to include 57 additional genes with FDR>=0.05 but nominal P-value<0.05. These additional genes include: *Apoe* and *Bin1. Bin1* encodes the bridging integrator 1, also known as amphiphysin 2, is the second most important risk locus for late onset Alzheimer’s disease, after apolipoprotein E (*Apoe*) [60, 61]. The two genes, *Apoe* and *Bin1*, rank 25th and 61th in the list of a total of 75 genes (18 genes with FDR<0.05 followed by 57 genes with nominal P-value<0.05). Both play a role in the *negative regulation of amyloid precursor protein catabolic process* (GO:1902992) and *tau protein binding* (GO:0048156). The Enrichr analysis [32, 33] reported following top GO terms: *regulation of neuron projection development* (GO:0010975; *Stmn2, Crmp1, Apoe, Cx3cl1, mark2), positive regulation of cell projection* organization (GO:0031346; *Stmn2, Crmp1, Apoe, Cx3cl1, mark2), phosphatidylserine metabolic process* (GO:0006658; *Lpcat4, Serinc1, Serinc5*), and *protein acylation* (GO:0043543; *Ing4, Abhd17b, Zdhhc17), potassium channel activity* (GO:0005267; *Kcnt2, Kcnh7, Kcnma1, Grik1*) and *methylation-dependent protein binding* (GO:0140034; *Ing4, Cbx5, Zmynd8*). The pre-ranked GSEA analysis [25] shows that regulatory changes are associated with *integrin signaling pathway, serotonin HTR1 group and FOS pathway*, and *glutamate neurotransmitter release cycle* (**Fig. 7D** inserts).

## Discussion

We present scTenifoldNet, a robust, unsupervised machine learning workflow that streamlines comparative GRN analyses with data from scRNAseq. The key feature of scTenifoldNet is to apply comparative network analysis with scRNAseq data. It detects differences in the cell population’s state between two samples in a sensitive and scalable manner. It provides the function of differential regulation (DR) analysis, which can be used to reveal subtle regulatory shifts of genes.

Today, differential expression (DE) analysis is still the primary method for the purpose of comparative analysis between scRNAseq samples (see, e.g., [27, 52, 62]). As scRNAseq data sets are becoming widely available, there will be more and more interest in comparing between samples. The scTenifoldNet-based DR analysis is expected to be adapted in more scenarios wherever DE analysis is applicable. scTenifoldNet learns and contrasts high-dimensional features of genes in scGRNs by examining global interactions between the genes. scTenifoldNet is more suitable for comparing highly similar samples, such as two populations of cells of the same type. scTenifoldNet is built as a robust, sensitive tool that can capture signals that are even confined to rare cell types.

To achieve technical requirements, we overcome several analytical barriers in developing scTenifoldNet. First, constructing scGRN from scRNAseq data, which consists of cells in many different states, is challenging at present. It is also difficult to control for technical noise in the data. To address these issues, we let scTenifoldNet begin with random cell subsampling. It is worthy noting that random cell subsampling can not only help dealing with the problem of cell heterogeneity, additional information of cells can be incorporated into subsampling schema. More specifically, in addition to the random subsampling using jackknife and bootstrap methods, we can adapt a semi-random subsampling schema, if cells in an input matrix are sorted according to pseudotime [63]. These cells can be subsampled using a pseudotime-guided method, with which sorted cells are sampled along the pseudotime trajectory. In such a way, the subsamples contain pseudotime information, and the multilayer scGRN constructed from these subsamples will contain the pseudotime-series information. In machine learning, many multilayer network analysis algorithms have been proposed [64–66]. With our pseudotime-series scGRN data, these algorithms will be relevant and applicable. Second, regulatory relationships between genes from scRNAseq data are difficult to establish, even though the data may theoretically capture a complete picture of the regulatory gene landscape. We consider principal component regression to stand out as a crucial method of building scGRNs. Principal component regression significantly outperforms the other GRN construction algorithms in all aspects of methodology metrics, including specificity, sensitivity, computational efficiency, and the required minimum number of cells. Importantly, principal component regression explicitly projects thousands of gene expression measurements into a low dimensional space to capture much of the observed variation. Principal component regression, therefore, establishes the relationship for each pair of genes after controlling for the most important background interactions. Third, in scTenifoldNet, the tensor denoising procedure effectively smooths edge weights across all networks in multilayer scGRNs. Third, scTenifoldNet performs nonlinear manifold alignment to align two networks. As such, two networks can be contrasted directly, and DR genes could be detected using distance in new coordinates of data in a low-dimensional space.

We validate the power of scTenifoldNet using real data sets coming from various studies and demonstrate that scTenifoldNet is sensitive to signals. Five real scRNAseq data sets are involved. These five data sets have one thing in common: they all have two sets of scRNAseq data—one from a treated group and the other from a control/untreated group. More importantly, in all five cases, we have sufficient prior knowledge about the biological system, from which the data is collected. Therefore, we have hypotheses about what transcriptional changes are expected to see before doing the analysis. For example, in the morphine response analysis, the causal factor of transcriptional responses, i.e., the morphine stimulus, is known and thus, we know what should be recovering through the analysis. Similarly, we had some clues in the examples of cetuximab and fibroblasts about what transcriptional changes we might be able to retrieve. By compiling all the findings from scTenifoldNet applications, we tested scTenifoldNet and showed that scTenifoldNet provides findings that are precise, specific and relevant to the biological systems and questions in the test. It is of significance to building a specific and sensitive tool like scTenifoldNet for the purpose of molecular mechanism studies using scRNAseq. This is because causal factors and their target genes remain unknown in many biological systems studied. If this is the case, it is crucial to apply the sensitive approach like scTenifoldNet, which may be in addition to the DE analysis, to unveil more gene candidates. Only then will we be able to scrutinize identified genes further to learn the mechanisms behind their actions in the whole system. We face such a challenge in many studies from unknown factors that cause the disorder. It is therefore critical that we adopt tools such as scTenifoldNet, instead of relying solely on conventional DE analysis, to tackle this big data analysis problem.

In summary, scRNAseq enables the study of cellular, molecular components, and dynamics of complex biological systems at single-cell resolution. To unravel the regulatory mechanisms underlying cell behaviors, novel computational methods are essential for understanding the complexity in scRNAseq data (e.g., scGRNs) that surpasses human interpretative ability. We anticipate that, when applied to real scRNAseq data, our machine learning workflow implemented in scTenifoldNet, can help achieve breakthroughs by deciphering the full cellular and molecular complexity of the data through constructing and comparing scGRNs.

## Methods

The scTenifoldNet workflow takes two scRNAseq expression matrices as inputs. The two matrices are supposed to be obtained from the same type of cells of two samples, such as those of different treatments or from diseased and healthy subjects. The purpose of the analysis is to identify genes whose transcriptional regulation is shifted between the two samples. The whole workflow consists of five steps: cell subsampling, network construction, network denoising, manifold alignment, and module detection.

### Cell subsampling

Instead of using all cells of each sample to construct a single GRN, we randomly subsample cells multiple times to obtain a set of subsampled cell populations. This subsampling strategy is to ensure the robustness of results against cell heterogeneity in samples. Subsampling of each sample is performed as follows: assuming the sample has *M* cells, *m* cells (*m* < *M*) are randomly selected to form a subsampled cell population. The process is repeated with cell replacement for *t* times to produce a set of *t* subsampled cell populations.

### Network construction

For a given expression matrix, a principal component regression-based network construction method [5] is adopted to construct scGRN. Principal component regression is a popular multiple regression method, where the original explanatory variables are the first subject to a principal component analysis (PCA) and then the response variable is regressed on the few leading principal components. By regressing on *M* principal components (*M* << *n*, where *n* is the total number of genes in the expression matrix), principal component regression mitigates the overfitting and reduces the computation time. To build an scGRN, each time we focus on one gene (referred to as the target gene) and apply the principal component regression method, treating the expression level of the target gene as the response variable and the expression levels of other genes as the explanatory variables. The regression coefficients from principal component regression are then used to measure the strength of the association of the target gene and other genes and to construct the scGRN. We repeat this process *n* times, each time with one gene as the target gene. At the end, the interaction strengths between all possible gene pairs are obtained and an adjacency matrix is formed. The details of applying the principal component regression method to a scRNAseq expression data matrix are described as follows.

More specifically, suppose 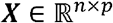 is the gene expression matrix with *n* genes and *p* cells. The *i*^th^ row of ***X***, denoted by 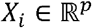 represents the gene expression level of the *i*^th^ gene in the *p* cells. We construct a data matrix 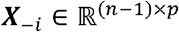 by deleting *X_i_* from ***X***. To estimate the effects of the other *n* – 1 genes to the *i*^th^ gene, we build a principal component regression model for *X_i_*. First, we apply PCA to 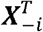, and take the first *M* leading principal components to construct 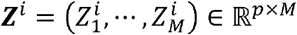 where 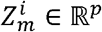 is the *m^th^* principal component of 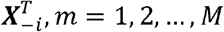. Mathematically, 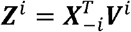, where 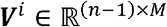 is the PC loading matrix for the first *M* leading principal components, satisfying 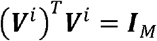. Secondly, the principal component regression method regresses ***X**_i_* on ***Z**^i^* and solves the following optimization problem:

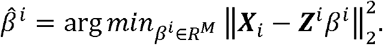

Then, 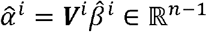 quantifies the effects of the other *n* – 1 genes to the *i*^th^ gene. After performing principal component regression on each gene, we collect 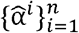 together and construct an *n* × *n* weighted adjacency matrix ***W*** of the gene-gene interaction network. The *i*^th^ row of ***W*** is 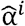, and the diagonal entries of ***W*** are all 0. Then we retain interactions with top *α*% (=5% by default) absolute value in the matrix to obtain the scGRN adjacency matrix.

### Tensor decomposition

For each of the *t* subsamples of cells obtained in the cell subsampling step, we construct a network using principal component regression, as described above. Each network is represented as a *n* × *n* adjacency matrix; the adjacency matrices of the *t* networks can be stacked to form a third-order tensor 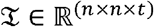. To remove the noise in the adjacency matrices and extract important latent factors, the CANDECOMP/PARAFAC (CP) tensor decomposition is applied. Similar to the truncated singular value decomposition (SVD) of a matrix, the CP decomposition approximates the tensor by a summation of multiple rank-one tensors [11]. More specifically, for our problem:

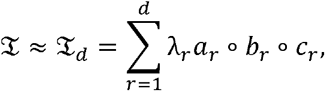

where ∘ denotes the outer product, 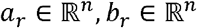, and 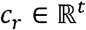 are unit-norm vectors, and *λ_r_* is a scalar. In the CP decomposition, 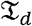 is the denoised tensor of 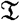, which assumes that the valid information of 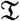 can be described by *d* rank-one tensors, and the rest part 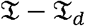 is mostly noise.

We use the function `cp` in the R package ‘rTensor’ to do the CP decomposition. For each sample, the reconstructed tensor 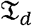 includes *t* denoised scGRNs. We then calculate the average of associated *t* denoised networks to obtain the overall stable network. We further normalize entries by dividing them by their maximum absolute value to obtain the final scGRNs for the given sample. For later use, denote the denoised adjacency matrices for the two samples as 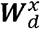 and 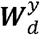.

### Manifold alignment

After obtaining 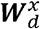 and 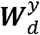, we compare them to identify the regulatory changes and associated genes and modules. Instead of directly comparing the two *n*×*n* adjacency matrices, we apply manifold alignment to build comparable low-dimensional features and compare these features of genes between two samples, while maintaining the structural information of the two scGRNs [19]. Manifold alignment is used here to match the local and no-linear structures among the data points of 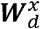 and 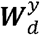 and project them to the same low-dimensional space. Specifically, we use 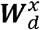 and 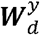 to denote the pairwise similarity matrices obtained by applying the principal component regression-based network construction method, and then denoising through tensor decomposition on the two initial expression matrices, ***X*** and ***Y***. These similarity matrices serve as the input for manifold alignment to find the low-dimensional projections 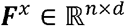 and 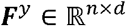 of genes from each sample, where *d* ≪ *n*. In terms of the underlying matrix representation, we use 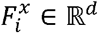 and 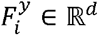 to denote the *i*^th^ row of ***F**^x^* and ***F**^y^* that reflect the features of the *i*^th^ gene in ***X*** and ***Y***, respectively.

We note that 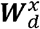 and 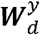 may include negative values, which means genes are negatively correlated. When the similarity matrix contains negative edge weights, the properties of the corresponding Laplacian are not entirely well understood [67]. We propose two methods to deal with this problem. The first method is to directly take the absolute value of 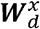 and 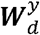 as the similarity matrices, in which we regard that the highly-negative correlated genes also support a strong functional relationship. In the second method, we add 1 to all entries in 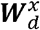 and 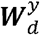, transforming the range of 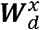 and 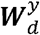 from [−1,1] to [0,2]. As a result, all original negative relationships have a transformed value in and all original positive relationships have a transformed value in (1,2]. In this case, the projected features of two genes with a positive correlation will be closer than those with a negative correlation. For convenience, we still use 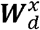 and 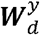 to denote the transformed similarity matrices of two data sets.

Now we propose a specific manifold alignment method to find appropriate low-dimensional projections of each gene. Our manifold alignment should trade off the following two requirements: (1) the projections of the same *i*^th^ gene in two samples should be relatively close in the projected space; and (2) if *i*^th^ gene and *j*^th^ gene in sample 1 are functionally related, their projections 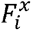 and 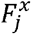 should be close in the projected space, and the same is true for sample 2. We minimize the following loss function:

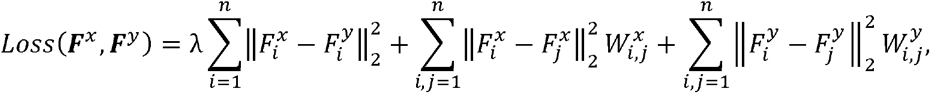

where 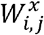 and 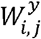 denote the (*i*, *j*) entry of 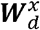 and 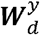 respectively. The first term of the loss function requires the similarity between corresponding genes across two samples; the second and third terms are regularizers preserving the local similarity of genes in each of the two networks. λ is an allocation parameter to balance the effects of two requirements.

One way to minimize the loss function is by using an algorithm similar to Laplacian eigenmaps [68], which requires the adjacency matrix to be symmetry, but in our case both 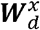 and 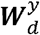 are asymmetric. Notice that if we symmetrize 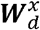 and 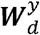 by 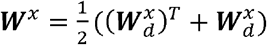 and 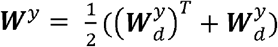, and again denote 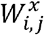 and 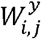 as the (*i*, *j*) entry of ***W**^x^* and ***W**^y^*, then the value of the loss function won’t be changed. Thus, minimizing the loss function based on the symmetrized adjacency matrices, ***W**^x^* and ***W**^y^*, is equivalent to using the original adjacency matrices, 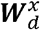 and 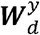. Based on this observation, using linear algebra, we can write the loss function into the matrix form as *Loss*(***F**^x^*, ***F**^y^*) = 2*trace*(***F**^T^**LF***), where: 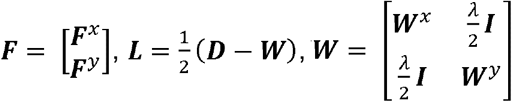, and ***D*** is a diagonal matrix with *D_ii_* = ∑*_i_ W_ij_*. ***L*** is called a graph Laplacian matrix. The default selection of *λ* is 0.9 times the mean value of the row sums of ***W**^x^* and ***W**^y^*. By further adding the constraint ***F**^T^**F*** = ***I*** to remove the arbitrary scaling factor, minimizing *Loss*(***F**^x^*, ***F**^y^*) is equivalent to solving an eigenvalue problem. The solution for ***F*** = [*f*_1_, *f*_2_, …, *f_d_*] is given by *d* eigenvectors corresponding to the *d* smallest nonzero eigenvalues of ***L*** [69].

### Determination of p-value of DR genes

With 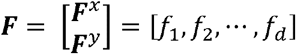 obtained in manifold alignment, we calculate the distance *d_j_* between projected data points of two samples for each gene. One may declare significant genes according to the ranking of *d_j_*’s. To avoid arbitrariness in deciding the number of selected genes, we propose to use the Chi-square distribution to determine significant genes. Specifically, 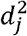 is derived from the summation of squares of the differences of projected representations of gene *j* for two samples, whose distribution could be approximately Chi-square. To adjust the scale of the distribution, we compute the scaled fold-change defined as 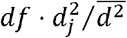 for each gene *j*, where 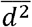 denotes the average of 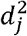 among all the tested genes. The scaled fold-change approximately follows the Chi-square distribution with the degree of freedom *df* if the gene does not perform differently in the two samples. By using the upper tail (*P*[*X* > *x*]) of the Chi-square distribution, we assign the *P*-values for genes and adjust them for multiple testing using the Benjamini-Hochberg (B-H) FDR correction [70]. To determine *df*, since the number of the significant genes will increase as *df* increases, we use *df* = 1 to make a conservative selection of genes with high precision.

### Functional enrichment analyses

Functional enrichment analysis of gene sets was performed using Enrichr [32, 33], which is a web-based, integrative enrichment analysis application based on more than 100 curated gene set libraries. The test of enriched TF targets was performed using the ChIP-X enrichment analysis (ChEA) [34] based on comprehensive results from ChIP-seq studies. Finally, predefined gene sets from the REACTOME, BioPlanet and KEGG databases were tested for the enriched functions using the pre-ranked Gene Set Enrichment Analysis (GSEA) [25].

### Simulations of scRNAseq data and benchmarking of network methods

To test the performance of our workflow, we generated synthetic data sets using SERGIO, a single-cell expression simulator guided by gene regulatory networks (GRNs) [71]. SERGIO allows for the simulation of scRNAseq data while considering the linear and nonlinear influences of regulatory interactions between genes. SERGIO takes a user-provided GRN to define the interactions and generates expression profiles of genes in steady-state using systems of stochastic differential equations derived from the chemical Langevin equation. The time-course of mRNA concentration of gene *i* is modeled by:

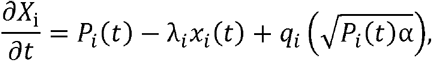

where *x_i_* is the expression of gene *i, P_i_* is its production rate, which reflects the influence of its regulators as identified by the given GRN, λ_*i*_ is the decay rate, *q_i_* is the noise amplitude in the transcription of gene *i*, and *α* is an independent Gaussian white noise process. In order to obtain the mRNA concentrations as a function of time, the above stochastic differential equation is integrated for all genes as follows:

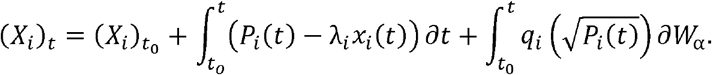

The simulation was focused on testing and comparing the performance of principal component regression and several other methods (SCC, MI, GENIE3) using sparse data without imputation. The relationships between 100 genes were simulated as they belong to two major modules containing 40 and 60 genes, respectively. Each module is under the influence of one TF. We used the steady-state simulations to synthesize data to generate expression profiles of 100 genes, according to the parameter setting for two modules.

For each one of the tested methods, we randomly select *n* = {10, 50, 100, 500, 1000, 2000, 3000} cells from the simulated data for ten times and build ten scGRN. For each *n*, relevance measurements (accuracy and recall) were evaluated for each of the ten networks using the match of the sign of the relationships between genes to compute the following formulas: *Accuracy* = (*TP* + *TN*)/(*TP* + *TN* + *FP* + *FN*), and *Recall* = *TP*/(*TP* + *FN*), where *T, P, F*, and *N* stands for true, positive, false and negative, respectively. For the MI and GENIE3 methods that only provide positive values, the median value was used as the center point and then the values were scaled to [−1,1] by dividing them over the maximum absolute value.

### Code availability

scTenifoldNet has been implemented in R. The source code is available at https://github.com/cailab-tamu/scTenifoldNet, which also includes the code of the benchmarking method, auxiliary functions, and example datasets (including the simulated data used to generate **Fig. 2**). The scTenifoldNet R package is available at the CRAN repository https://cran.r-project.org/web/packages/scTenifoldNet/.

## Acknowledgments

This research was funded by Texas A&M University 2019 X-Grants and the NIH grant R21AI126219 for J.J.C.

